# Fatty acid amide hydrolase drives adult mammary gland development by promoting luminal cell differentiation

**DOI:** 10.1101/2023.04.03.535417

**Authors:** Isabel Tundidor, Marta Seijo-Vila, Sandra Blasco-Benito, María Rubert-Hernández, Gema Moreno-Bueno, Laura Bindila, Rubén Fernández de la Rosa, Manuel Guzmán, Cristina Sánchez, Eduardo Pérez-Gómez

## Abstract

Mammary gland development occurs primarily in adulthood, undergoing extensive expansion during puberty followed by cycles of functional specialization and regression with every round of pregnancy/lactation/involution. This process is ultimately driven by the coordinated proliferation and differentiation of mammary epithelial cells. However, the endogenous molecular factors regulating these developmental dynamics are still poorly defined. Endocannabinoid signaling is known to determine cell fate-related events during the development of different organs in the central nervous system and the periphery. Here, we report that the endocannabinoid-degrading enzyme fatty acid amide hydrolase (FAAH) plays a pivotal role in adult mammary gland development. Specifically, it is required for luminal lineage specification in the mammary gland, and it promotes hormone-driven secretory differentiation of mammary epithelial cells by controlling the endogenous levels of anandamide and the subsequent activation of cannabinoid CB_1_ receptors. Together, our findings shed light on the role of the endocannabinoid system in breast development and point to FAAH as a therapeutic target in milk-production deficits.

## Introduction

The mammary gland is a unique glandular organ that distinguishes mammals from all other animals and whose paramount function is to produce and secrete milk to nourish the offspring. Architecturally, the virgin mammary gland consists of a bilayered ductal tree with a luminal layer that contains hormone receptor-positive (HR+) cells and hormone receptor-negative (HR-) cells primed for milk production (alveolar progenitors), as well as an outer basal layer that contains mostly contractile myoepithelial cells, but also bipotent mammary stem cells (MaSCs) (Visvader and Stingl 2014). During puberty, steroid hormones drive extensive expansion of the ductal system to fill the mammary fat pad and, when pregnancy occurs, the influence of prolactin orchestrates the secretory differentiation of alveolar progenitors and the development of the lobuloalveolar units specialized in milk production. After weaning, the epithelial compartment undergoes massive cell death and returns to a morphological pre-pregnancy state in a process known as post-lactational regression (Watson and Khaled 2020). Determining the differentiation dynamics that drive these postnatal developmental stages is not only essential to understand normal mammary gland function but also the etiology of pathological events as breast cancer.

The endocannabinoid system (ECS) is a cell-communication system that plays multiple roles in the regulation of cell physiology. This system is principally composed of (i) the G protein-coupled, cannabinoid-selective receptors CB_1_R and CB_2_R, (ii) their endogenous ligands (the endocannabinoids) anandamide (*N*-arachidonoylethanolamine, AEA) and 2-arachidonoylglycerol (2-AG), and (iii) the enzymes that synthesize and metabolize these ligands (Fowler 2021). Previous studies have shown that endocannabinoid-associated signaling pathways control cell fate decisions such as proliferation, differentiation, and survival, especially in the central nervous system (Galve-Roperh et al. 2013). Here, we aimed to decipher the possible role of the ECS in adult mammary gland development by characterizing and modulating the expression of the AEA-degrading enzyme fatty acid amide hydrolase (FAAH), a key molecular determinant of the endogenous AEA tone. We found that the expression of this enzyme is deeply associated to the luminal lineage in the adult mammary gland, and that it acts as a driver of lactogenic differentiation *in vitro* (by downregulating the endocannabinoid tone and its activity at CB_1_R) and *in vivo* (it is necessary for the generation of hormone-sensing luminal cells in the mammary gland). Taken together, our findings unveil the key relevance of the ECS in the cellular dynamics that occur during human lactation, and may provide further insights into the interplay between mammary gland development and breast cancer.

## Results

### FAAH expression is associated with luminally-differentiated and hormone receptor-positive mammary epithelial cells

In order to investigate the role of FAAH in the adult mammary gland development, we first assessed its mRNA levels and those of other related elements of the ECS in human mammary epithelial cell (MEC) populations through publicly available datasets. Traditionally, cell sorting techniques for specific cell surface markers have been used to prospectively isolate MEC subpopulations from mammary tissue. More specifically, the combination of antibodies against epithelial cell adhesion molecule (EpCAM) (or the heat stable antigen CD24 as surrogate) and integrin α6 (CD49f) allows segregation between the differentiated luminal subset (EpCAM^+^ CD49f^-^), the luminal progenitor subset (EpCAM^+^ CD49f^+^), and the basal/MaSC subset (EpCAM^-^ CD49f^+^), which is thought to contain both differentiated basal cells and bipotent MaSCs (Stingl et al. 2006). Microarray analyses on FACS-sorted populations purified from reduction mammoplasties (Kannan et al. 2013) revealed that *FAAH* mRNA levels were significantly higher in the differentiated luminal subset than in the basal/MaSC subset, while expression levels in the luminal progenitor subset fell between the other two (Fig. 1a). Similar data were obtained after the analysis of two additional studies involving gene expression profiling of human MEC subpopulations ((Raouf et al. 2008);(Lim et al. 2009)) (Figs. 1b, c). One of these studies (Raouf et al. 2008) provided further segregation of the basal/MaSC subset into mature basal (CD49f^−^ MUC1/CD133^-^ CD10/THY1^+^) and bipotent progenitor (CD49f^+^ MUC1/CD133^-^ CD10/THY1^+^) subsets. No differences in *FAAH* levels were observed in these two subpopulations (Fig. 1c). To validate these findings at the protein level, we performed immunofluorescence staining of FAAH in human adult mammary gland samples. Consistent with our previous observations, FAAH colocalized exclusively with luminal cells in the inner layer of the inactive lactiferous duct [characterized by the expression of cytokeratin 8 (CK8)], but not with basal cells in the outer layer [characterized by the expression of smooth muscle actin (SMA)] (Fig. 1d). Together, these results suggest that FAAH expression is associated with the luminal lineage in the adult mammary gland.

**Figure 1.**
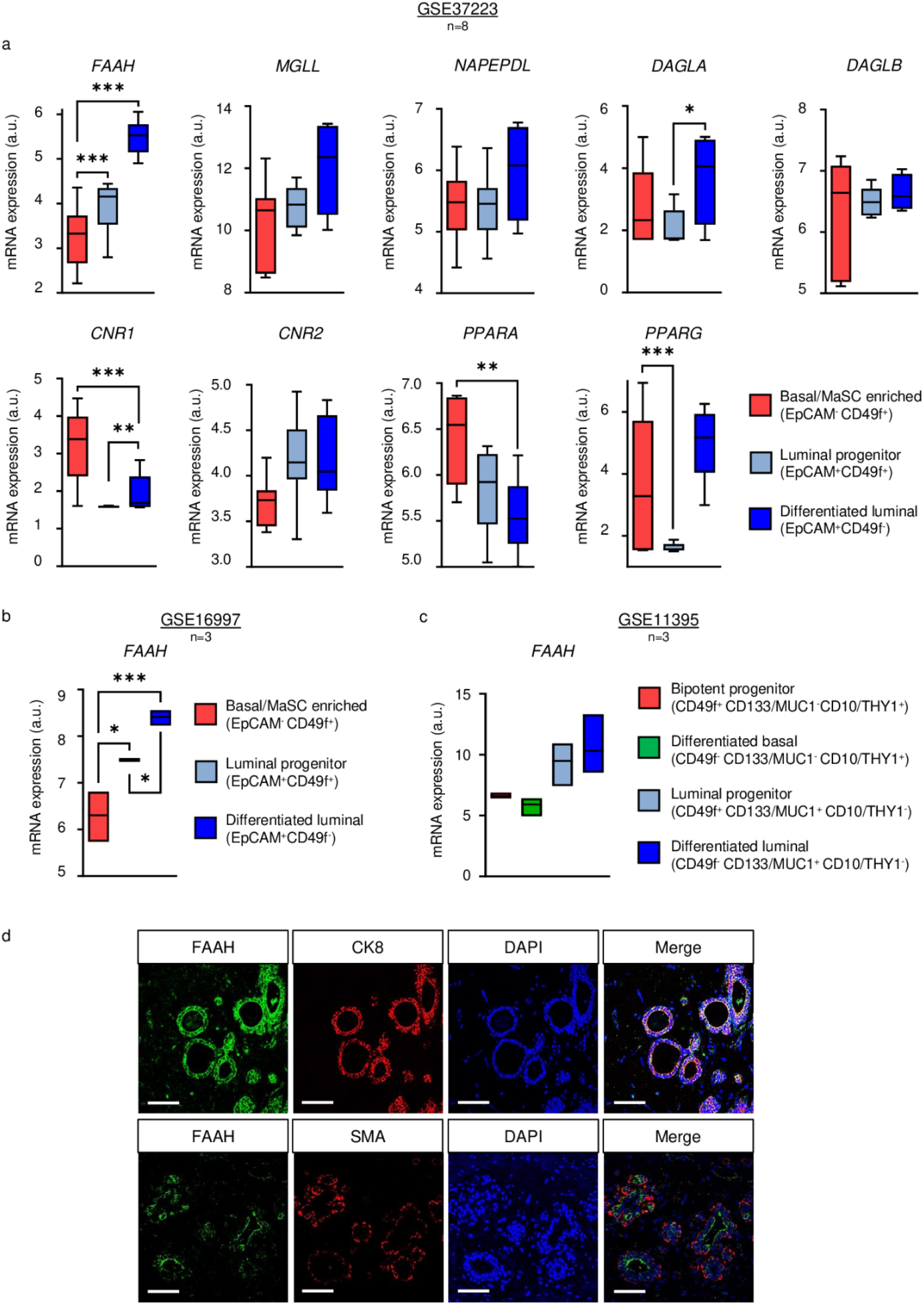
Expression of FAAH and related elements of the ECS in human mammary gland cell populations. **a-c** Data were obtained from the microarray dataset published by (Kannan et al. 2013) (a), (Lim et al. 2009) (b), and (Raouf et al. 2008) (c). Data are presented as mean values ± SEM of n=8 (a) and n=3 (b,c) biologically independent samples. *FAAH*: fatty acid amide hydrolase; *MGL*: monoacylglycerol lipase; *NAPEPLD*: N-acyl phosphatidylethanolamine-specific phospholipase D; *DAGLA*, *DAGLB*: diacylglycerol lipase A and B; *CNR1*: cannabinoid receptor 1; *CNR2*: cannabinoid receptor 2; *PPARA*, *PPARG*: peroxisome proliferator activated receptor alpha and gamma. ANOVA: * p < 0.05, ** p < 0.01, *** p < 0.001. **d** Representative immunofluorescence staining of FAAH with cytokeratin 8 (CK8) as a luminal marker (upper panels) and with smooth muscle actin (SMA) as a basal marker (lower panels) in human mammary samples obtained from breast reductions. Cell nuclei are stained in blue. Scale bar = 50 μm.

No remarkable differences between MEC populations were observed for the mRNA levels of other endocannabinoid (eCB)-synthesizing (*NAPEPLD*, *DAGLA,* and *DAGLB*) or eCB-degrading (*MGLL*) enzymes (Fig. 1a). Regarding cannabinoid receptors CB_1_R and CB_2_R (encoded by *CNR1* and *CNR2*, respectively) and other cannabinoid-related receptors that may be potentially engaged and activated by FAAH substrates (such as the peroxisome proliferator-activated receptors PPARα and PPARγ, encoded by *PPARA* and *PPARG*, respectively), their mRNA levels also presented differential expression between cell subsets, with *CNR1* and *PPARA* transcripts showing opposite patterns to that of FAAH, *i.e.,* higher in the basal/MaSC subset (Fig. 1a).

In recent years, single cell RNA sequencing (scRNA-seq) has allowed to comprehensively address the heterogeneity of the mammary epithelium as well as the identification of functionally relevant subpopulations within the luminal and basal compartments. To better characterize the expression pattern of FAAH in the mammary epithelial cell hierarchy, we next studied its mRNA levels in a dataset containing scRNA-seq profiling of mouse mammary glands from four adult developmental time points: nulliparous, mid gestation, lactation, and post involution (Bach et al. 2017) (Fig. 2a). Paralleling our observations in human tissue, *Faah* transcripts were predominantly expressed in cell clusters of differentiated luminal cells (C3, C4, C5, C8 and C11) and, to a lesser extent, in cell clusters of luminal progenitors (C1, C2, C6, C7, C9 and C10), while it was absent from basal (C12-C14) and MaSC (*i.e.*, Procr^+^ cells, C15) clusters (Fig. 2a). Importantly, the highest expression of *Faah* mRNA within luminal differentiated cells was found in C3 and C4 clusters, which showed characteristics of hormone-sensing cells such as high expression levels of genes encoding for hormone (estrogen, progesterone, and prolactin) receptors (*Esr1*, *Pgr*, and *Prlr*, respectively), while its expression was lower in HR-alveolar cells (C8 and C11), characterized by the expression of β-casein (*Csn2*) (Bach et al. 2017) (Fig. 2a). Similar observations were made in human tissue after the analysis of *FAAH* mRNA levels in the UCSC Genome Browser, which integrates 24 K human mammary epithelial cell atlases obtained by scRNA-seq (Saeki et al. 2021) (Fig. 2b), as well as in the Human Protein Atlas (Thul et al. 2017) (Supplementary Fig. 1).

**Figure 2.**
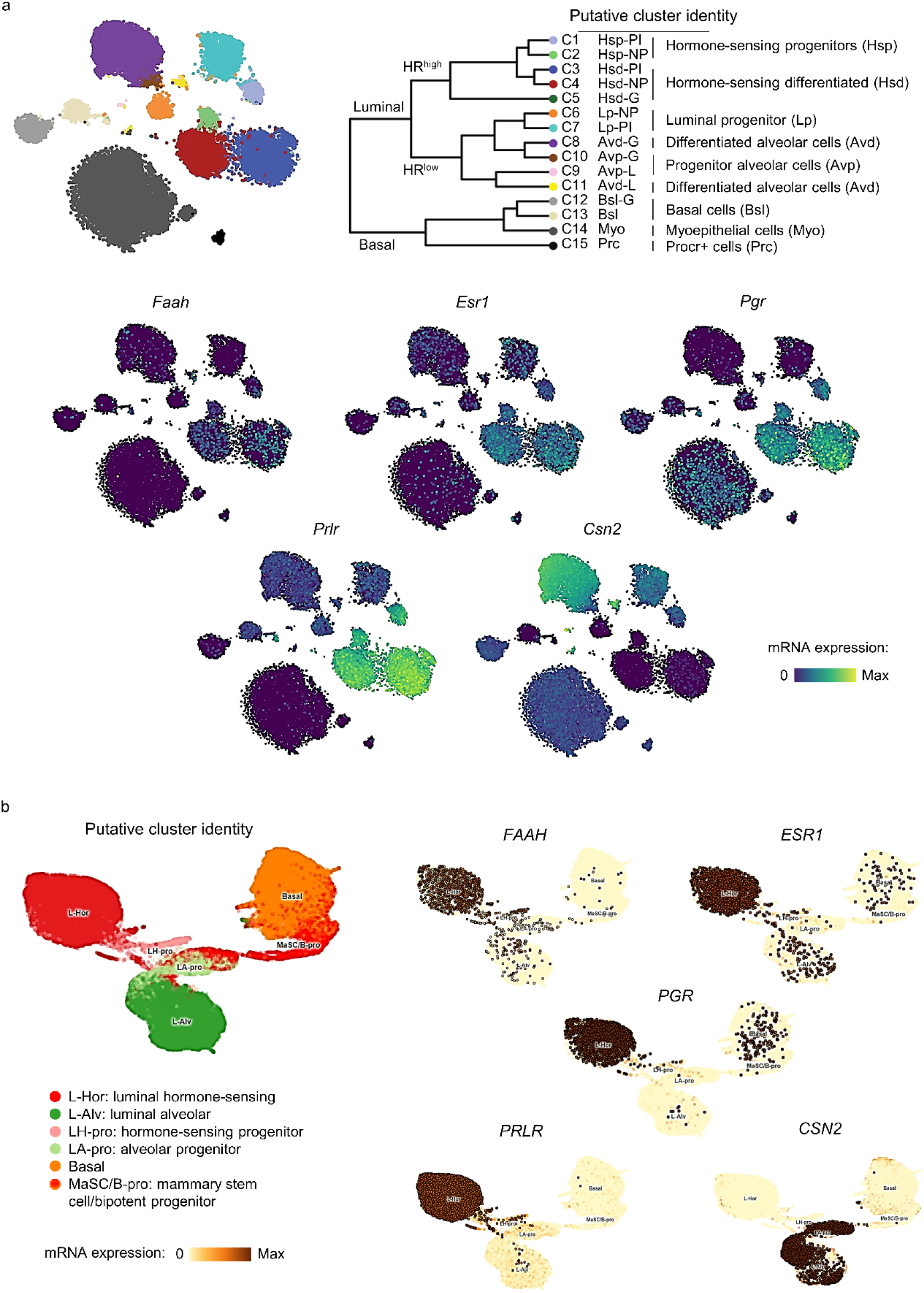
FAAH is expressed in luminal hormone-sensing cell populations in all stages of adult mammary gland development. **a,b** t-SNEs plots representing mRNA expression of *FAAH* and cell population-specific genes in putative cell clusters of mammary cell populations defined by scRNA-seq analysis of developing mouse mammary glands as published by Bach *et al*., 2017 (a) and Saeki *et al*., 2021 (b). t-SNEs plots are colored by the normalized log-transformed expression of each of the genes. *ESR1*: estrogen receptor alpha; *PGR*: progesterone receptor; *PRLR*: prolactin receptor; *CSN2*: β-Casein.

Together, these results show a strong association between high FAAH expression and differentiated, HR+ cell phenotypes within the luminal lineage, and thus suggest that FAAH may be involved in cell differentiation and lineage specification in the mammary gland.

### FAAH drives mammary epithelial cell differentiation *in vitro*

To validate the hypothesis that FAAH is involved in the differentiation of MECs, we next investigated its role in an *in vitro* model of lactogenic differentiation, a crucial process that occurs when pregnancy is advanced to expand the ductal system and differentiate it to generate milk secretion sites. Specifically, we used HC11 cells, a murine MEC model that displays progenitor-like features and retains its ability to differentiate into milk-producing cells in response to the addition of lactogenic hormones prolactin (PRL) and dexamethasone (DXM) (Ball et al. 1988). First, we differentiated HC11 cells from their progenitor-like (*i.e.*, undifferentiated) state to an intermediate or “competent” (*i.e*., predifferentiated) state (by removing EGF from the culture medium), from which the addition of the lactogenic stimuli triggers their final differentiation into milk-producing (*i.e*., differentiated) cells (Fig. 3a). Differentiation was monitored by following the gradual increase in β-casein expression, which is a widely accepted differentiation marker in this cell line. In line with our previous observations, HC11 cell differentiation (*i.e*., β-casein increase) was accompanied by a concomitant increase in FAAH expression (Fig. 3b).

**Figure 3.**
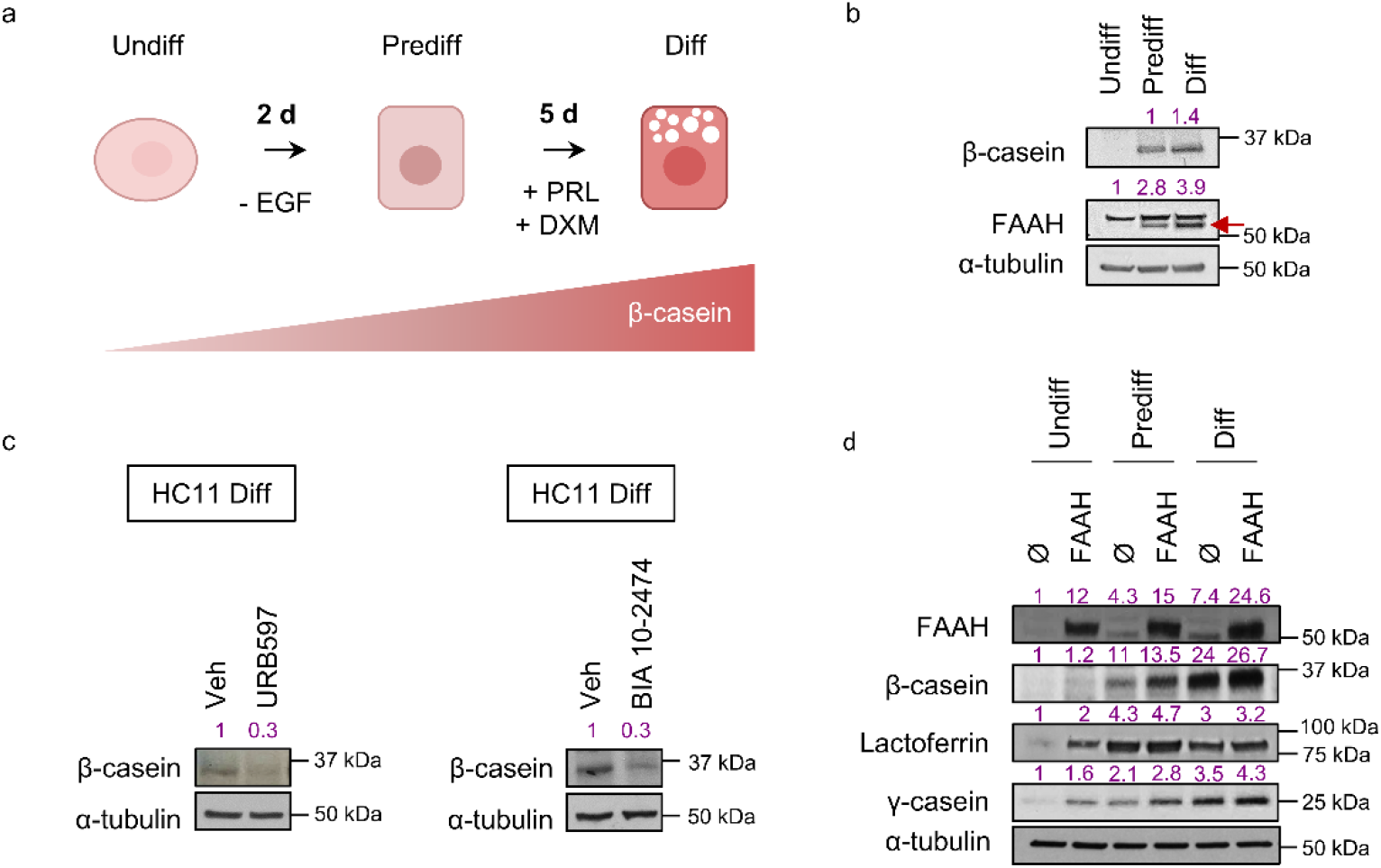
FAAH drives lactogenic differentiation of MECs *in vitro*. **a** Schematic representation of the differentiation protocol of HC11 cells**. b** Representative WB analysis of β-casein and FAAH at the three differentiation points. The specific band for FAAH in HC11 cells (pointed with an arrow) was determined by WB analysis of FAAH^-/-^ mouse tissue (Supplementary Fig. 2a). **c** Representative WB analysis of β-casein in HC11 cells that have completed the differentiation protocol in the presence of the FAAH inhibitors URB597 or BIA 10-2494 at 5 μM. **d** Representative WB analysis of β-casein and other milk proteins during the complete differentiation of HC11 cells harboring an empty (Ø) or a FAAH overexpression vector. Densitometric values for all WBs after normalization against α-tubulin and control condition are depicted in purple.

To try to establish a cause-effect link between the increase in FAAH expression and HC11 differentiation, we next blocked FAAH activity during the differentiation process by using the selective enzyme inhibitor URB597. FAAH inhibition led to an impaired lactogenic differentiation, as determined by the prevention of the increase in β-casein at the final stage of the process (Fig. 3c). These data were mimicked by using a different FAAH inhibitor (BIA 10-2474) (Fig. 3c), indicating that the catalytic activity of FAAH is required for the differentiation of HC11 cells.

To further confirm that FAAH is involved in the differentiation process associated with the lactogenic switch, we overexpressed the enzyme in HC11 cells via a lentiviral delivery system. As shown in Fig. 3d, FAAH overexpression led to an increase in β-casein and other milk proteins corresponding to molecular weights previously observed for lactoferrin and γ-Casein (with apparent molecular weights of 80 and 25 kDa, respectively) (Boumahrou et al. 2011). This occurred even in those stages where lactogenic hormones had not been added yet (*i.e*., undifferentiated and predifferentiated), which further supports the notion that FAAH drives the differentiation of mammary epithelial cells.

### FAAH drives MEC differentiation by decreasing anandamide-mediated activation of cannabinoid receptors

The data obtained upon modulating the activity and expression of FAAH support the idea that the enzymatic activity of FAAH participates in the control of lactogenic differentiation of MECs *in vitro*, and therefore, they put forward the possible involvement of one or several FAAH substrates in this process. FAAH is the main enzyme that terminates the action of AEA and other bioactive lipids of the *N*-acylethanolamine (NAE) family, such as palmitoylethanolamide (PEA) and oleoylethanolamide (OEA), that lack affinity for CB_1_R and CB_2_R but are active at proposed non-canonical cannabinoid receptors as PPARα and PPARγ. To unmask the primary molecular target of the inhibition of lactogenic differentiation that may be driven by FAAH substrates, we blocked a battery of receptors that can be engaged and activated by these compounds. Differentiating HC11 cells were incubated with selective antagonists for CB_1_R (SR141716, herein referred to as SR1), CB_2_R (SR144528, herein referred to as SR2), PPARα (GW 6471) or PPARγ (GW 9662) in the presence of URB597. Only SR1 completely prevented the downregulation of β-casein triggered by URB597 (Fig. 4a), indicating that CB_1_R mediates the FAAH-evoked inhibition of MEC differentiation. A partial prevention of the downregulation of β-casein by SR2 was also observed, suggesting CB_2_R might be partly involved too (Fig. 4a).

**Figure 4.**
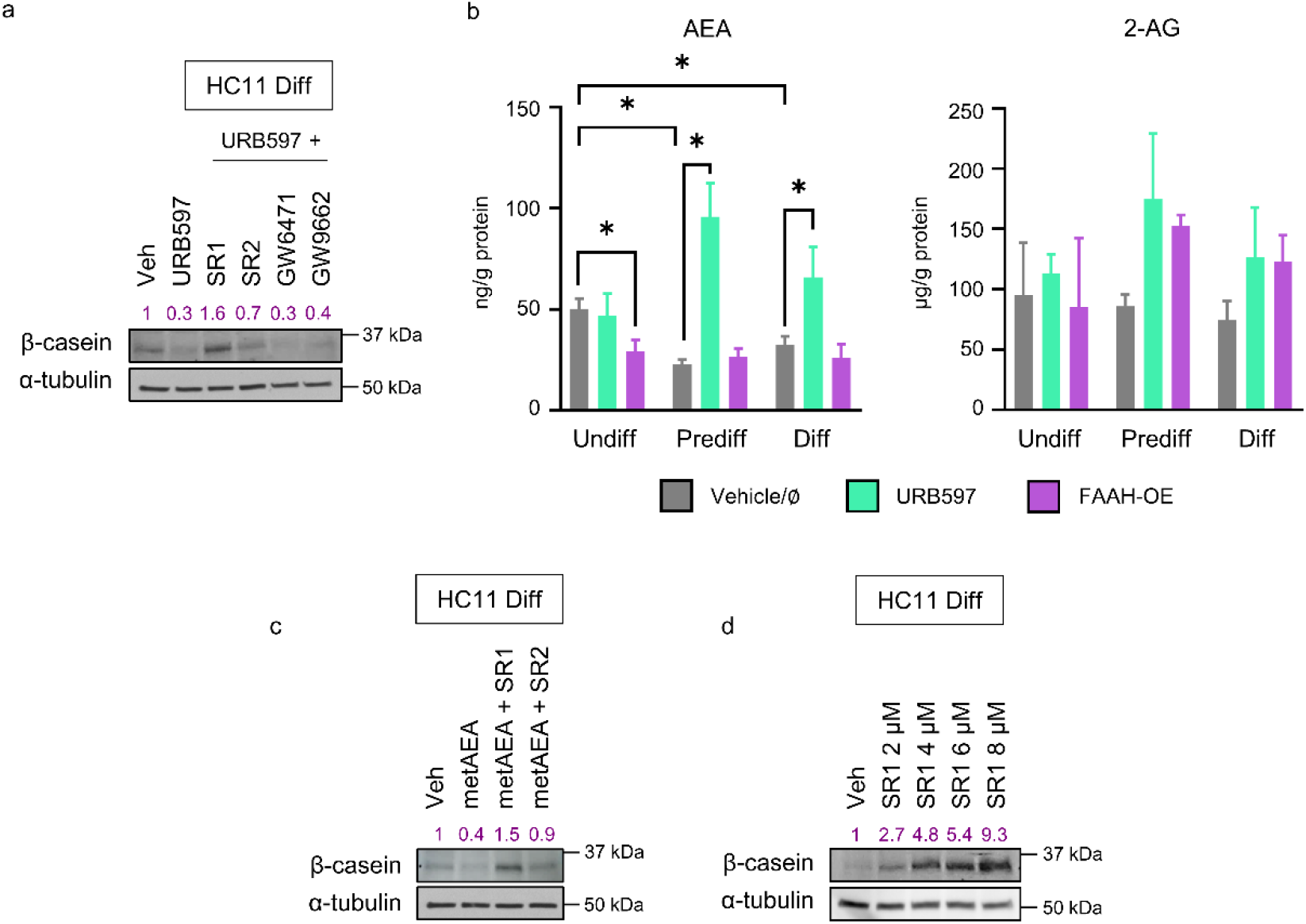
FAAH triggers HC11 cell differentiation by decreasing AEA-mediated cannabinoid receptor activity. **a** Representative WB analysis of β-casein in HC11 cells after being differentiated in the presence of URB597 at 5 μM and selective receptor antagonists at 1 μM. **b** Levels of the endocannabinoids anandamide (AEA) and 2- arachidonoylglycerol (2-AG) in FAAH-modulated HC11 cells normalized to protein content. Values represent mean ± SEM of n=3 biologically independent samples. Student’s t-test: * p < 0.05. **c** Representative WB analysis of β-casein in HC11 cells after being differentiated in the presence of metAEA at 0.5 μM and selective antagonists to CB_1_R and CB_2_R at 1 μM. **d** Representative WB analysis of β-casein in HC11 cells after being differentiated in the presence of increasing concentrations of the CB_1_R selective antagonist SR1. Densitometric values after normalization against α-tubulin and control condition are depicted in purple.

Given that only CB_1_R and CB_2_R (but not PPARs) were shown to mediate the effect of FAAH on MEC differentiation, we next analyzed the cellular levels of endocannabinoids in the HC11 *in vitro* differentiation model. We first observed that, in parallel to the increase in FAAH expression, AEA levels decreased throughout the differentiation process, while no changes in 2-AG levels were found (Fig. 4b). As expected, FAAH inhibition by URB597 led to an increase in AEA levels in the three different differentiation stages (Fig. 4b), while the opposite approach (*i.e*., FAAH overexpression) led to a decrease in AEA levels in undifferentiated, but not in predifferentiated and differentiated HC11 cells, probably due to a “ceiling effect” owing to the already high FAAH levels expressed by the cells in those stages (Fig. 4b).

All the aforementioned observations point to an activation of CB_1_R by endogenous AEA as the responsible for the blockade of cell differentiation found upon FAAH inhibition. In order to further study this causative link, we challenged differentiating HC11 cells with methanandamide (metAEA), a non-hydrolysable AEA analogue. MetAEA phenocopied the effect of FAAH pharmacological inhibition (*i.e*., it impaired HC11 differentiation), and this effect was prevented by both SR1 and SR2 (Fig. 4c), further reinforcing that the AEA tone, as driven by FAAH, is involved in the control of MEC differentiation through the activation of cannabinoid receptors. Of interest, SR1 not only prevented URB597 (Fig. 4a) and metAEA (Fig. 4c) effect on β-casein levels but increased them above those of the vehicle-treated cells. To better understand the importance of this observation, we treated HC11 cells during differentiation with SR1 alone and detected a concentration-dependent increase in β-casein (Fig. 4d), *i.e.*, an induction of cell differentiation. Together, these data demonstrate that FAAH drives MEC differentiation by decreasing AEA-mediated activation of CB_1_R.

### FAAH expression is regulated by estradiol in mammary epithelial cells

Next, we aimed to find upstream regulators of FAAH expression that help clarify the biological significance of its role in MEC differentiation. Hormones are the main regulators of the adult phase of mammary gland development. While estrogen and progesterone are key for ductal growth and tertiary branching during puberty and early pregnancy, PRL and glucocorticoids mostly drive alveolar morphogenesis that culminates at lactation (Neville et al. 2002). Of note, data in Fig. 2 and Supplementary Fig. 1 showed a high FAAH expression in hormone receptor-positive luminal cells. To determine whether there is a functional link between these hormones and FAAH activity, we treated undifferentiated HC11 cells with increasing concentrations of estradiol (E_2_), progesterone (P_4_), PRL and DXM. Among all treatments, only E_2_ was able to stimulate FAAH expression (Fig. 5a). This effect was maximum 2 h after E_2_ addition and quickly dissipated afterwards (Fig. 5b). The connection between estrogen signaling and FAAH was further strengthen by the analysis of mouse mammary gland *Faah* mRNA levels during pregnancy, lactation, and involution (Clarkson et al. 2004), which showed a parallel variation with that of the estrogen receptor (*Esr1*) (Fig. 5c). Together, these data raise the possibility that the regulation of FAAH expression in MECs by E_2_ has physiological relevance for the accomplishment of mammary gland development during the adult phase.

**Figure 5.**
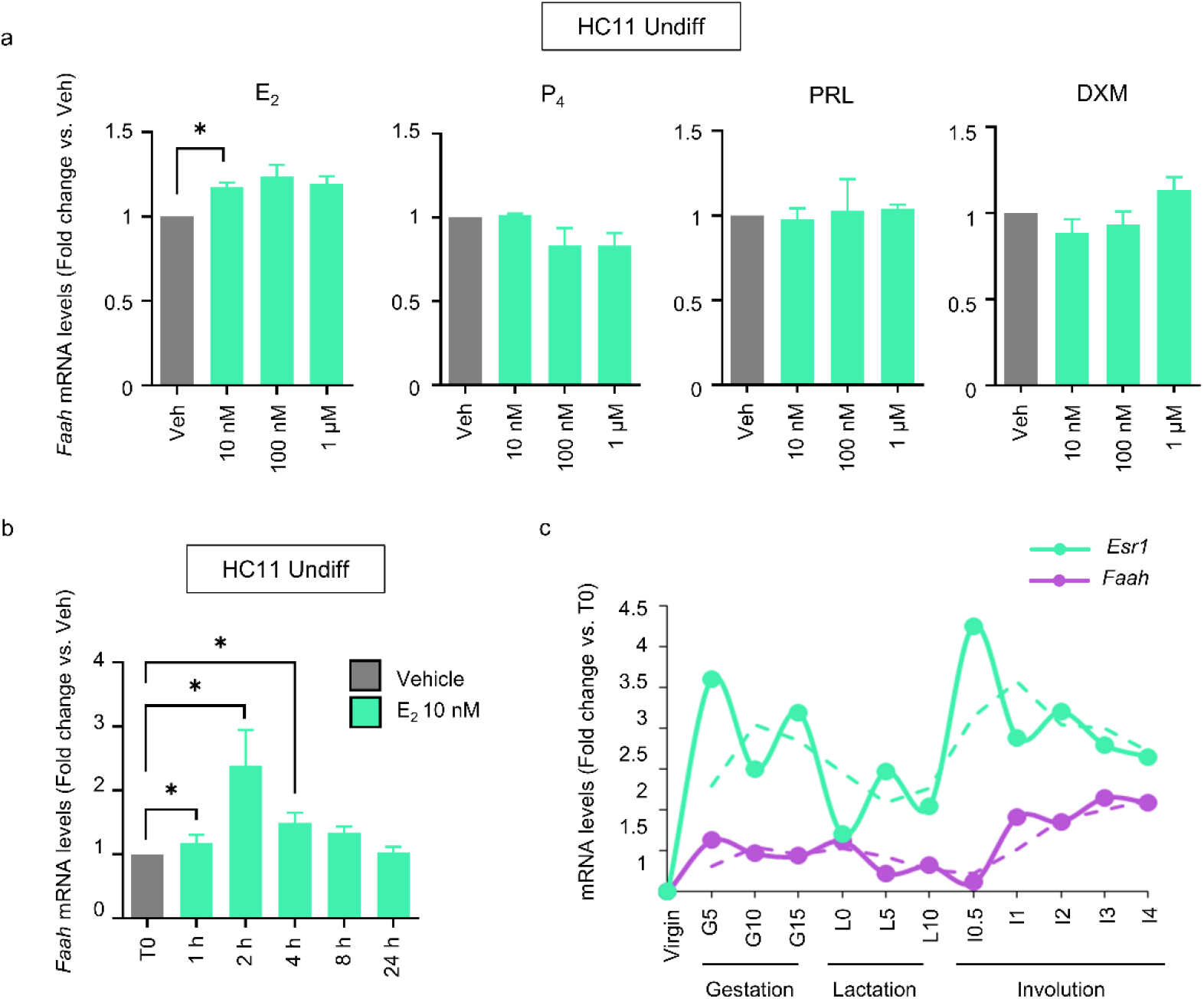
FAAH expression is regulated by estradiol in undifferentiated HC11 cells. **a, b** qPCR analysis of *Faah* mRNA expression in undifferentiated HC11 cells after treatment with increasing concentrations of estradiol (E_2_), progesterone (P_4_), prolactin (PRL) and dexamethasone (DXM) for 4 h (a) or with 10 nM E_2_ for the indicated times (b). Expression levels were normalized against *Tbp*. Values represent mean ± SEM of n=3 (a) and n=4 (b) biologically independent samples. **c** *Faah* and *Esr1* mRNA expression at different time points of adult mammary gland development, obtained from the microarray dataset published by (Clarkson et al. 2004).

### FAAH is necessary for the generation of hormone-sensing luminal cells and a functional PRLR-β-casein axis in the mouse mammary gland

As FAAH was found to be a relevant driver of MEC differentiation *in vitro*, we hypothesized that its genetic inactivation would have deleterious effects over mammary gland development in adult mice. Whole-mount morphological and histological analyses of mouse tissue showed that the structure of FAAH ^-/-^ mammary glands at different adult developmental points (virgin, gestation, lactation, and involution) was essentially normal compared to that of FAAH ^+/+^ littermates, and that knocking out FAAH did not affect the architecture or the number of mammary ducts or alveoli (Fig. 6a). However, flow cytometry-based analysis of Sca1 and CD49b markers (which allows to segregate the CD24^+^ CD49f^-^ luminal compartment into hormone-sensing and alveolar cells) revealed that virgin FAAH ^-/-^ mammary glands contained a significantly lower proportion of hormone-sensing cells (and a consequent higher proportion of alveolar cells) in their luminal layer compared to FAAH ^+/+^ controls (Fig. 6b), thus suggesting that knocking out FAAH may have a negative impact on how MECs sense and integrate hormonal cues in order to drive mammary gland development. Given that the developmental features mainly orchestrated by E_2_ and P_4_ (*i.e.*, duct length and tertiary branching during puberty and gestation) seemed unaffected in FAAH ^-/-^ mice (Fig. 6a), and based on the role that FAAH had demonstrated to play in PRL-dependent secretory differentiation *in vitro* (Fig. 4), we hypothesized that FAAH ^-/-^ mammary glands may have functional defects during lactation, which is the developmental stage most dependent on PRL signaling. In line with this, Western blot analysis demonstrated that FAAH ^-/-^ mammary glands had lower expression levels of β-casein at different time points of lactation, and this was accompanied by a downregulation of PRL receptor (PRLR) at the same time points (Fig. 6c). Interestingly, treatment of HC11 cells with the FAAH inhibitor URB597 also downregulated mRNA levels of *Prlr* (Fig. 6d). All this suggests that FAAH ^-/-^ mammary glands may not undergo complete lactogenic differentiation because they exhibit partial resistance to PRL action. In line with this notion, no differences on PRL expression levels were found between FAAH ^+/+^ and FAAH ^-/-^ mammary glands during lactation (Fig. 6e).

**Figure 6.**
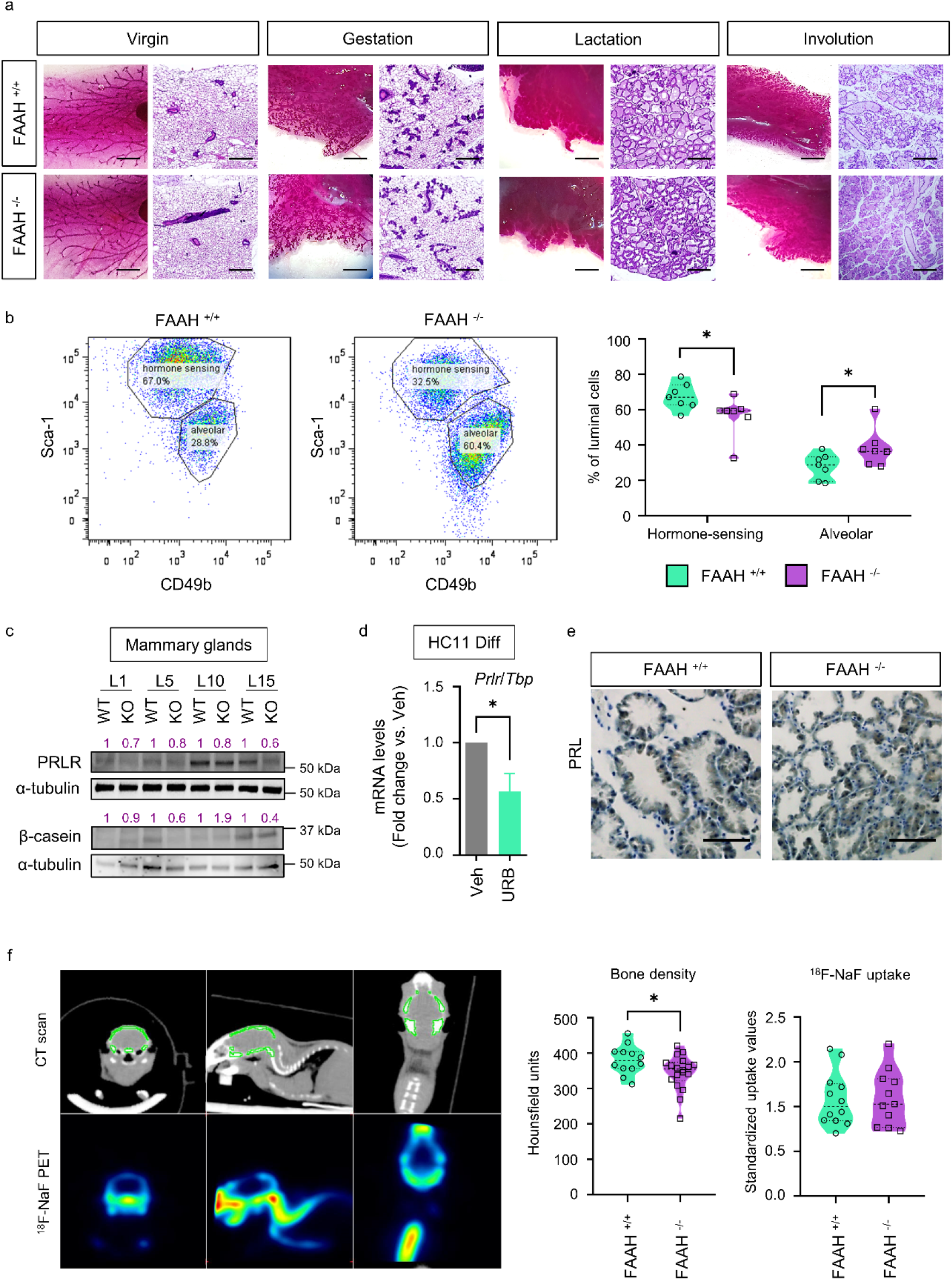
FAAH is necessary for the generation of hormone-sensing luminal cells and a functional PRLR-β-casein axis in the mouse mammary gland. **a** Representative whole-mount pictures and H&E-stained sections of FAAH ^+/+^ and FAAH ^-/-^ mouse mammary glands harvested at different time points of adult development (virgin, gestation day 15, lactation day 15, and involution day 2). n = 2 mice per time point. Scale bar = 1 mm (whole mount) and 200 μm (H&E). **b** Representative flow cytometric analysis of MEC populations of FAAH ^+/+^ and FAAH ^-/-^ virgin mammary glands. CD24^+^ CD49f^-^ luminal cells were isolated and then separated into hormone-sensing or alveolar lineages based on their expression of Sca-1 and CD49b (see Methods). n = 7 mice per group. **c** Representative WB analysis of β-casein and PRLR at L1, L5, L10 and L15. Densitometric values after normalization against α-tubulin and WT condition for every time point are depicted in purple. **d** qPCR analysis of *Prlr* mRNA expression in HC11 cells after being differentiated in the presence of the FAAH inhibitor URB597 at 5 μM. Expression levels were normalized against *Tbp*. Data are shown as mean ± SEM of n=7 biologically independent experiments. Student’s t-test: * p < 0.05. **e** IHC detection of PRL in samples at L10. Scale bar = 200 μm. **f** Left, representative images of the computed tomography (CT), showing the area of study (the skull) in green, and of the positron emission tomography (PET) with ^18^F-NaF, showing a heat map visualization of the ^18^F-NaF uptake. Right, results of the CT scan (skull bone density) and the ^18^F-NaF PET (^18^F-NaF uptake) expressed in Hounsfield units and standardized uptake values, respectively. n = 12 and 18 mice in the FAAH ^+/+^ and FAAH ^-/-^ breastfeeding group, respectively.

We next aimed to identify the functional consequences of the abnormal cell composition of the FAAH ^-/-^ mammary glands. As mentioned above, we hypothesized that pups from females lacking FAAH may have lactation-related deficits. First, we analyzed the body weight of pups fed by FAAH ^-/-^ females and found no differences when compared with those fed by FAAH ^+/+^ animals. Next, and given that FAAH ^-/-^ mice had lower β-casein levels in their mammary glands, we hypothesized that pups that were breastfed by FAAH^-/-^ females may present defects in their bone density. Casein phosphopeptides (derived from caseins by tryptic digestion) are pivotal milk components for infant health owing to their ability to bind and solubilize calcium ions, improving calcium absorption at the intestinal level (Sun et al. 2018). FAAH ^+/+^ and FAAH ^-/-^ female mice were mated with studs of the opposite genotype (so that the litters of both FAAH ^+/+^ and FAAH ^-/-^ were the same genotype, FAAH ^+/-^) and allowed to breastfeed for a 4-week period, after which pups were weaned and subjected to a computed axial tomography (CT) to assess bone deposition up to that point. CT scans revealed that pups that had been breastfed by FAAH^-/-^ females had significantly lower bone density compared to those breastfed by FAAH ^+/+^ females (Fig. 6f). To determine their osteogenic activity, pups were subjected to positron emission tomography (PET) with ^18^F-NaF right after the CT scan and no differences were found between the two experimental groups (Fig. 6f), pointing to a nutritionally-deficient breast milk composition of FAAH ^-/-^ females (rather than a metabolic defect) as the primary cause for these differences.

## Discussion

The reproductive phase of mammary gland development is a hormone-dependent morphogenetic process in which the different epithelial cell types collaborate to generate lobuloalveolar units specialized in milk production. HR+ (or hormone-sensing) luminal cells play a key role in this process as they sense systemic hormonal changes and transduce them into paracrine instructions for neighboring HR- (or alveolar progenitors) cells, which, in response, will differentiate into alveolar cells able to synthesize and secrete milk (Visvader and Stingl 2014). In this study, we demonstrate that the AEA- degrading enzyme FAAH is highly expressed by the hormone-sensing lineage of the adult mammary gland, and that it is involved in the acquisition of the hormone-sensing phenotype that integrates hormonal signaling in order to drive MEC differentiation.

Here, we used the HC11 cell line as a model of progenitor-like cells with the capability to differentiate into milk-producing cells in response to the addition of lactogenic stimuli. Treatment of these cells with the FAAH inhibitor URB597 blocked lactogenic differentiation in a cannabinoid receptor-evoked manner, and this was mimicked by the AEA analogue metAEA, thus pointing to an elevated endogenous AEA tone triggered by FAAH inhibition as the main responsible for the phenotype in the former case. Over the years, attention has focused on the possible role of AEA and other endocannabinoids in regulating cell differentiation, which might account for some physio-pathological effects of these lipid mediators (Galve-Roperh et al. 2013). For example, AEA inhibits the differentiation of human keratinocytes (Paradisi et al. 2008) and neurons (Rueda et al. 2002) through CB_1_R-mediated attenuation of cell type-specific differentiation pathways. Lactogenic failure in our FAAH-deficient models (both *in vitro* and *in vivo*) was accompanied by a downregulation of PRLR expression levels, pointing to an involvement of the PRLR pathway in the observed phenotypic alterations. However, metAEA treatment of HC11 cells was unable to do so (Supplementary Fig. 2b). In spite of this, AEA has been previously reported to mediate CB_1_R-dependent downregulation of PRLR in breast cancer cells -thus rendering them insensitive to the mitogenic action of PRL (De Petrocellis et al. 1998). Therefore, a possible regulation by AEA of the PRLR pathway also in MECs should not be discarded. Future studies may inquire further into the relationship between the signaling pathways downstream CB_1_R activation and those involved in MEC differentiation to clarify the role of FAAH in this process.

Of note, treatment of differentiating HC11 cells with the CB_1_R-selective antagonist SR141716 (SR1) was sufficient to enhance cell-fate specification in a concentration-dependent fashion, thus pointing to either a ligand-independent constitutive activation of CB_1_R or a basal endocannabinoid tone acting on CB_1_R and actively inhibiting cell differentiation in MECs. Although the former cannot be ruled out, our results support the latter hypothesis, given that constitutive activation of FAAH (accompanied by downregulation of its mains substrates) mimicked CB_1_R antagonism, *i.e*., it promoted HC11 differentiation. Together, these observations support the existence of an AEA- driven, basal CB_1_R activity that prevents differentiation in luminal cells and is ultimately controlled by FAAH activity.

However, the contribution to inhibition of lactogenic differentiation of (i) FAAH substrates other than AEA that lack affinity for CB_1_R, and (ii) other endogenous CB_1_R agonists whose levels are not directly regulated by FAAH, cannot be ruled out. Regarding the first scenario, it is well-known that PEA and OEA, two FAAH substrates, can downregulate the expression and activity of FAAH in breast cancer cells (Bisogno et al. 1998; Di Marzo et al. 2001). It could be argued that, by downregulating FAAH, these could further enhance the anti-differentiation effects of AEA in MECs. Regarding the second possibility, it is worth noticing that the levels of 2-AG in HC11 cells also suffered a notable (although not statistically significant) increase in response to URB597. Although monoacylglycerol lipase (MAGL) is responsible for 85% of 2-AG hydrolysis in the mouse brain, it can also be cleaved into glycerol and arachidonic acid by other hydrolases such as FAAH (Maccarrone et al. 2015). However, 2-AG levels did not decrease upon FAAH constitutive activation, so we concluded that most likely it does not contribute to the FAAH-controlled endocannabinoid tone acting at CB_1_R in HC11 cells.

Our data also demonstrate that transgenic mice lacking FAAH show an aberrant development of the mammary gland during lactation, suggesting that FAAH may be necessary for this process to occur *in vivo*. In this regard, several hormones involved in the development of the mammary gland have been found to regulate *FAAH* gene expression through direct or indirect interaction with its promoter. Research in human neuroblastoma cells first identified estrogen and glucocorticoid response elements (EREs and GREs, respectively) in the *FAAH* promoter that regulated transcriptional activity independent of their ligand (Waleh et al. 2002). In the uterus, *FAAH* expression is downregulated by sex hormones (MacCarrone et al. 2000), and the opposite has been reported in mouse Sertoli cells, in which E_2_ induces ER binding to ERE2/3 sites in the *Faah* promoter and activates its transcription (Grimaldi et al. 2012). Likewise, there is evidence that progesterone binds and regulates transcriptional activity of the *FAAH* gene in human T lymphocytes through the transcription factor Ikaros (Maccarrone et al. 2003). Here, we demonstrate that *FAAH* expression is induced by E_2_ in MECs, and put forward the biological importance of ER-mediated regulation of FAAH in mammary gland development.

The evidence gathered in this study reveals the profound importance of FAAH in the differentiation dynamics of MECs in the adult mammary gland, and raises potentially important implications for the pathogenesis of breast cancer. Recent work from our lab established that FAAH genetic inactivation in the MMTV-neu transgenic mouse model preceded the formation of highly proliferative, more aggressive tumors (Tundidor et al. 2023). Here, we demonstrated that mammary glands from FAAH ^-/-^ mice show an expanded population of alveolar progenitors, which are the likely target for transformation in the MMTV-neu model (Henry et al. 2004). Therefore, it could be argued that, by regulating MEC plasticity in the normal mammary gland, FAAH could be involved in the early stages of HER2/neu-induced tumorigenesis, and probably other breast cancer subtypes.

Aside from the insight provided into the potential role of FAAH in breast cancer, our findings may have translational relevance on their own. Inadequate milk production affects many women after giving birth, but the molecular determinants underlying lactational failure are still poorly understood (Olsen and Gordon 1990; Kent et al. 2012). Recent studies demonstrated that treatment of HC11 cells with the plant-derived cannabinoids Δ^9^-tetrahydrocannabinol and cannabidiol reduced the levels of the main nutritional components of the milk, thus reiterating the risks of cannabis consumption during pregnancy and lactation (Josan et al. 2022). In the same line, here we demonstrate that FAAH ^-/-^ females (which are subjected to a higher endocannabinoid tone) show a significant decrease in β-casein levels, and that this phenotypic alteration translates into a functional deficiency during lactation, as the pups fed by FAAH ^-/-^ females showed lower bone density at weaning.

Taken together, these findings put forward the important implication of the ECS in adult mammary gland development, and therefore contribute to the identification of physiological barriers for appropriate milk production which are necessary to develop novel interventions to support breastfeeding. In addition, they represent a note of caution for the use of cannabis during pregnancy, which constitutes a topic of scientific and social debate.

## Methods

### Reagents

The FAAH inhibitor URB597 and the non-hydrolysable anandamide analog R-1 methanandamide (metAEA) were purchased from Cayman Chemical (Ann Arbor, MI, USA). The FAAH inhibitor BIA 10-2474 was purchased from Medchem Express (Monmouth Junction, NJ, USA). Selective antagonists of CB_1_R (SR141716, herein referred to as SR1), CB_2_R (SR144528, herein referred to as SR2), PPARα (GW 6471) and PPARγ (GW 9662) were purchased from Tocris Bioscience (Bristol, UK).

### HC11 lactogenic differentiation

The murine cell line HC11 was a kind donation by Dr. Castillo-Lluva (Complutense University, Madrid, Spain) and was authenticated by short tandem repeat profiling at the Genomics Core Facility of Instituto de Investigaciones Biomédicas Alberto Sols (Madrid, Spain). Cells were maintained in subconfluence in RPMI medium (Sigma-Aldrich, St. Louis, MO, USA) with 10% fetal bovine serum (FBS) (Gibco, Billings, MT, USA), 10 µg/mL insulin (Sigma-Aldrich), 20 ng/mL epidermal growth factor (EGF) (Gibco), 2 mM L-glutamine (Lonza, Basel, Switzerland), 10 mM HEPES (Lonza) and 1% penicillin/streptomycin (Lonza). To induce their differentiation, subconfluent HC11 cells were trypsinized and seeded in complete medium at a density of 8.5 x 10^5^ cells per well in 6-well plates. When they reached confluence (usually, overnight), complete medium was changed to EGF-free medium and predifferentiated cells were obtained after 48 h of growth in this medium. To obtain completely differentiated cells, predifferentiated cells were then treated for additional 5 days with differentiation medium: EGF-free medium containing 1 μg/mL prolactin (PRL) (Sigma-Aldrich) and 100 nM dexamethasone (DXM) (Sigma-Aldrich). Differentiation medium was changed and added fresh on day 3.

### Lentiviral transduction

For lentiviral transduction of HC11 cells, a second-generation packaging system was used. Commercial packaging (psPAX2) and envelope (pMD2.G) plasmids (Addgene, Watertown, MA, USA), as well as the pReceiver-Lv225 (Genecopoeia, Rockville, MD, USA), were kindly donated by Dr. Fernández-Piqueras (Centro de Biología Molecular Severo Ochoa, Madrid, Spain). EX-hFAAH-Lv225 transfer plasmid was home generated by subcloning human *FAAH* into the XhoI and BstbI sites of pReceiver-Lv225. The empty (Ø) vector EX-NEG-Lv225 was used as negative control. Briefly, lentiviruses were generated by transfecting 3 x 10^6^ HEK-293T cells in antibiotic-free medium with 6 µg of total DNA in a 1:2:3 ratio of pMD2.G:psPAX2:transfer plasmid using polyethylenimine (PEI) (Sigma-Aldrich). 48 h after transfection, conditioned medium containing recombinant lentiviruses was collected, filtered through 0.45 μm sterilization filters and added to HC11 cells together with 8 μg/mL polybrene (Sigma-Aldrich). Transduction efficiency was monitored by checking the expression of green fluorescent protein (GFP), which usually happened 48-72 h later. Antibiotic selection was not performed as the transfection efficiency was usually near 100% and lasted through the complete HC11 differentiation protocol.

### Endocannabinoid measurement

For sample preparation, cells were grown in 6-cm dishes and scraped off in ice-cold PBS. They were centrifuged twice at 2000 g for 10’ at 4 °C and pellets were frozen in liquid nitrogen and stored at −80 °C until processed. Lipid extraction and parallel quantification of endocannabinoids was performed at the lipidomics unit of the *Universitätsmedizin* associated to the University of Mainz (Germany) as previously described (Schwitter et al. 2023).

### RNA extraction and real-time quantitative PCR (Q-PCR)

RNA was isolated with Nucleozol™ (Macherey-Nagel, Düren, Germany) following the extraction protocol suggested by the manufacturer. Reverse transcription (RT) was performed from 2-3 µg of RNA with the Transcriptor First Strand cDNA Synthesis Kit (Roche Life Sciences, Basel, Switzerland) using random hexamer primers. Real-time quantitative RT-PCR (Q-PCR) was performed in 384-well plates using the LightCycler® Multiplex DNA Master (Roche Life Sciences) and hybridization probes from the Universal Probe Library Set (Roche Life Sciences). The relative expression ratio of target/reference gene was calculated with the `ΔΔ`Ct method (Pfaffl 2001). Primer sequences for m*FAAH* were GCA GGT GGG CTG TTC AGT (sense) and AAG CAG GGA TCC ACA AAG TC (antisense); for m*Prlr*, TCC CTG GTA TGG CAG ACT TT (sense) and AAC ATC TGC GAT GCT CAC CT (antisense); for m*Tbp*, GGG GAG CTG TGA TGT GAA GT (sense) and CCA GGA AAT AAT TCT GGC TCA (antisense).

### Protein extraction and Western blot (WB)

Cells were scraped off in RIPA buffer (0.1% SDS, 0.5% sodium deoxycholate, 1% NP40, 150 mM NaCl, 50 mM Tris–HCl pH 8.0, in PBS) supplemented with protease and phosphatase inhibitors (Sigma-Aldrich). 25 µg of total protein were boiled for 5’ at 95 °C, resolved by SDS-PAGE and transferred to PVDF membranes in a Trans-Blot® SD Semi-Dry Transfer Cell (Bio-Rad, Hercules, CA, USA). Unspecific binding to membranes was prevented by blocking in 5% bovine serum albumin (BSA) for 1 h at room temperature (RT). Primary antibodies were incubated overnight at 4 °C in the blocking solution. The following primary antibodies were used: anti-milk [Nordic-MUbio (Susteren, Netherlands) #RAM/TM, dilution 1:5000], anti-FAAH [Abcam (Cambridge, UK) #ab128917, dilution 1:1000], anti-CB_1_R [Frontier Institute (Nittobo Medical, Tokyo, Japan) #CB1-GP-Af530-1, dilution 1:500], anti-PRLR [Abcam #ab214303, dilution 1:1000], and anti-α-tubulin (Sigma-Aldrich #T9026, dilution 1:5000). Membranes were then washed three times with TBS-T, incubated for 1 h at RT with the corresponding HRP-conjugated secondary antibody (Cytiva Lifescience, Marlborough, MA, USA) and developed with self-prepared ECL reagent (1.25 mM luminol, 0.2 mM p-coumaric acid, 100 mM Tris–HCl pH 8.5, in dH_2_O). Densitometric analysis was performed with ImageJ™ software (Schindelin et al. 2012).

### Human samples

Human mammary gland sections from reduction mammoplasties were kindly given by Dr. Moreno-Bueno (MD Anderson Cancer Center, Madrid, Spain). All patients gave informed consent, and the study was authorized by the respective Hospital Ethics Committees.

### Immunofluorescence (IF)

Paraffin-embedded human tissue sections were de-waxed in isoparaffin H (Panreac, Castellar del Vallès, Spain) and hydrated by quick changes through serial ethanol baths. Antigen retrieval was performed by boiling in sodium citrate buffer (10 mM sodium citrate, 0.05% Tween 20, pH 6.0, in dH_2_O). Samples were subjected to several washes with dH_2_O and washing buffer (NaCl 8.3 g/L, Tris 1.2 g/L; pH 7.4). Blocking was performed with 10% goat serum in PBS, for 1 h at RT. Primary antibodies were diluted in Antibody Diluent (Dako) and incubated overnight at 4 °C. The following primary antibodies were used: anti-FAAH (Abcam #ab128917, dilution 1:100), anti-CK8 [DSHB (Iowa, IA, USA) #531826, dilution 1:50], and anti-SMA (DSHB #2289065, dilution 1:50) Samples were rinsed 2 x 5’ in PBS + 0.25% triton and incubated with fluorescent secondary antibodies [Invitrogen (Waltham, MA, USA), dilution 1:200] and 1 μg/mL DAPI for 1 h at RT. Finally, they were mounted with Mowiol® mounting medium. Fluorescence confocal images were acquired by using an Olympus FV1200 microscope.

### Animals

C57BL/6J FAAH ^-/-^ mice were kindly donated by Dr. Ben Cravatt’s laboratory (Scripps Research Institute, La Jolla, CA, USA). All procedures involving animals were performed with the approval of the Complutense University Animal Experimentation Committee and Madrid Regional Government according to the European official regulations. Animals were housed in the animal facility of the UCM School of Biology, under a 12 h light-dark cycle, and were allowed to feed and drink *ad libitum*.

### Mammary gland dissection

For studies on the virgin mammary gland, tissue was harvested from 12 to 18-week-old C57BL/6J FAAH ^+/+^ or FAAH ^-/-^ female mouse littermates. For studies during the reproductive phase of mammary gland development (gestation, lactation, and involution), 12 to 18-week-old C57BL/6J FAAH ^+/+^ or FAAH ^-/-^ female mice were mated with wild type studs and allowed to litter. For gestation studies, tissue was harvested at day 15 after vaginal plug detection. For lactation studies, tissue was harvested at days 1, 5, 10 and 15 postpartum. In this case, pups were removed from the cage 2 h before mammary gland extraction to make sure that the differences between FAAH ^+/+^ and FAAH ^-/-^ mice were not due to different sucking stimuli at the time of dissection. For involution studies, pups were force-weaned at 10 days of lactation, and tissue was harvested 2 days after. Integrity of pairs #1 and #5 was usually compromised after opening the skin and thus were never dissected. Pairs #2 and #3 were usually snap frozen and then used for RNA/protein extraction. Pair #4 was the preferred option for imaging studies as it was easier to remove integrally.

### Whole mounts and histology

For whole-mount analysis, mammary gland pair #4 was spread over a glass slide, soaked overnight in Carnoy’s fixative (glacial acetic acid:chloroform:absolut ethanol in a 1:3:6 ratio), hydrated through serial ethanol baths, stained with 0.5% Carmine Alum Stain (Sigma-Aldrich), dehydrated through serial ethanol baths and finally cleared in xylene before mounting in Eukitt® quick-hardening mounting medium (Sigma-Aldrich). For histology studies, mammary gland pair #4 was spread over a 6 cm dish, fixed overnight in 10% formalin, and transferred to 50% ethanol. Fixed mammary glands were then paraffin-embedded and sectioned at the Department of Medicine and Animal Surgery at UCM School of Veterinary, which also performed hematoxylin and eosin (H&E) staining. Prolactin IHC staining was performed in paraffin-embedded mouse tissue sections by the Histopathology Unit at the Spanish National Cancer (CNIO) (Madrid, Spain).

### Flow cytometry

Mammary gland pairs #2, #3 and #4 were pooled and processed to obtain a single cell suspension through mechanical disaggregation followed by 2 h of enzymatic digestion in DMEM + 125 μg/mL collagenase (Sigma-Aldrich), 0.25 µg/mL dispase (Sigma-Aldrich) and 10 µg/mL DNAse (Sigma-Aldrich) at 37 °C. Luminal cells were isolated through the staining with Alexa Fluor® 488 CD24 antibody [Biolegend (San Diego, CA, USA) #101816, dilution 1:1000] and Alexa Fluor® 647 CD49f antibody (Biolegend #313610, dilution 1:20) and subsequently segregated into hormone-sensing and alveolar through the staining with Brilliant Violet 605™ Sca-1 antibody (Biolegend #108133, dilution 1:40) and PE CD49b antibody (Biolegend #108907, dilution 1:80). In addition, cell suspension was stained for biotinylated CD31 [Thermo Fisher Scientific (Waltham, MA, USA) #13-0311-82, diluted 1:200], CD45 (Thermo Fisher Scientific #13-0451-82, diluted 1:400) and TER119 (Thermo Fisher Scientific #13-5921-82, diluted 1:100) + PE Streptavidin (Biolegend #405203, diluted 1:300) to exclude endothelial and hematopoietic lineages (Lin^+^ cells) from the analysis. Antibody incubations were sequentially performed in agitation for 20’ at 4 °C and protected from light. A washing step with FACS buffer (PBS + 1% FBS + 1% BSA + 0.02% sodium azide) was performed after each antibody incubation. Finally, cells were resuspended thoroughly in FACS buffer at a concentration of 1 x 10^6^ cells/mL and transferred to polystyrene round-bottom tubes. 7-aminoactinomycin D (7-AAD) (Biolegend) was used as death cell marker and added 15’ before the analysis. Data acquisition was performed in a FACSCalibur system (Becton Dickinson, Franklin Lakes, NJ, USA). Compensation was applied according to single-stained controls of the same cell type. For data analysis, FlowJo™ software was used. Unstained cells were used to set the blank for surface-marker expression after exclusion of cell debris, nonviable cells (7-AAD^-^), and Lin^+^ cells.

### Bone density and osteogenic activity

FAAH ^+/+^ and FAAH ^-/-^ female mice were mated with studs of the opposite genotype (FAAH ^-/-^ and FAAH ^+/+^, respectively) and allowed to litter. FAAH ^+/-^ pups were kept with their corresponding mother and breastfed for 4 weeks. To determine the degree of bone development achieved within this period (where their only food source was breastmilk), pups were weaned and their bone density analyzed by computed axial tomography (CT). Comparisons were made between pups fed by FAAH ^+/+^ and FAAH ^-/-^ mothers. To complement this information, and also right after weaning, the osteogenic activity of the pups was determined by positron emission tomography (PET) with ^18^F-Na fluoride (^18^F-NaF). The ^18^F-NaF tracer uptake indicates osteoblastic activity at sites of calcification, *i.e.*, rapidly growing or regenerating bone regions, and therefore provides metabolic rather than static information about the bone status. Both studies were performed with an Albira ARS II system (Bruker, Billerica, MA, USA). The CT scan was performed with a CT acquisition of 600 projections at 400 μA and 45 kV. For the PET examination, an intraperitoneal injection of ^18^F-NaF was made allowing the molecule to be incorporated for 60 minutes, after which a 30-minute static PET scan was performed. For the analysis of the CT images, the skull was used as the reference structure and the PMOD software v. 4.1 was used to create a segmentation of the volume (VOI or volume of interest) and to quantify its average density value, that was expressed in Hounsfield Units (HU). For the analysis of the PET images, numeric values were normalized to the animal weight and the injected amount of ^18^F-NaF, and were expressed as standardized uptake values (SUV). Both experimental groups were balanced in terms of the proportion of males and females and no differences by sex were found in any of the two parameters measured.

### Statistical analyses

Unpaired, independent groups of two were analyzed by 2-tailed Student’s t-test. For multi-group comparison, data were analyzed by one-way ANOVA with post Tukey’s multiple comparison test, or by two-way ANOVA when required. A p value of less than 0.05 was considered statistically significant. Unless otherwise stated, data are expressed as mean ± SEM from at least three biological replicates.

## Acknowledgements

We are indebted to Eva Resel for administrative support. We also want to thank the sample donors and the MD Anderson Cancer Center (Madrid, Spain) for the human specimens used in this study.

## Author contributions

I.T., C.S., and E.P-G. designed the research and wrote the manuscript. I.T., M.S-V., S.B-B., M.R-H., and R.F-R. performed the biological experiments. I.T. and E.P-G. performed computational and statistical analyses. I.T., C.S., M.G., and E.P-G. supervised the research and reviewed and revised the manuscript. L.B. and G.M-B. provided technical and material support.

