## Supplemental info for "Fatty acid amide hydrolase drives adult mammary gland development by promoting luminal cell differentiation"

Supplementary Figure 1

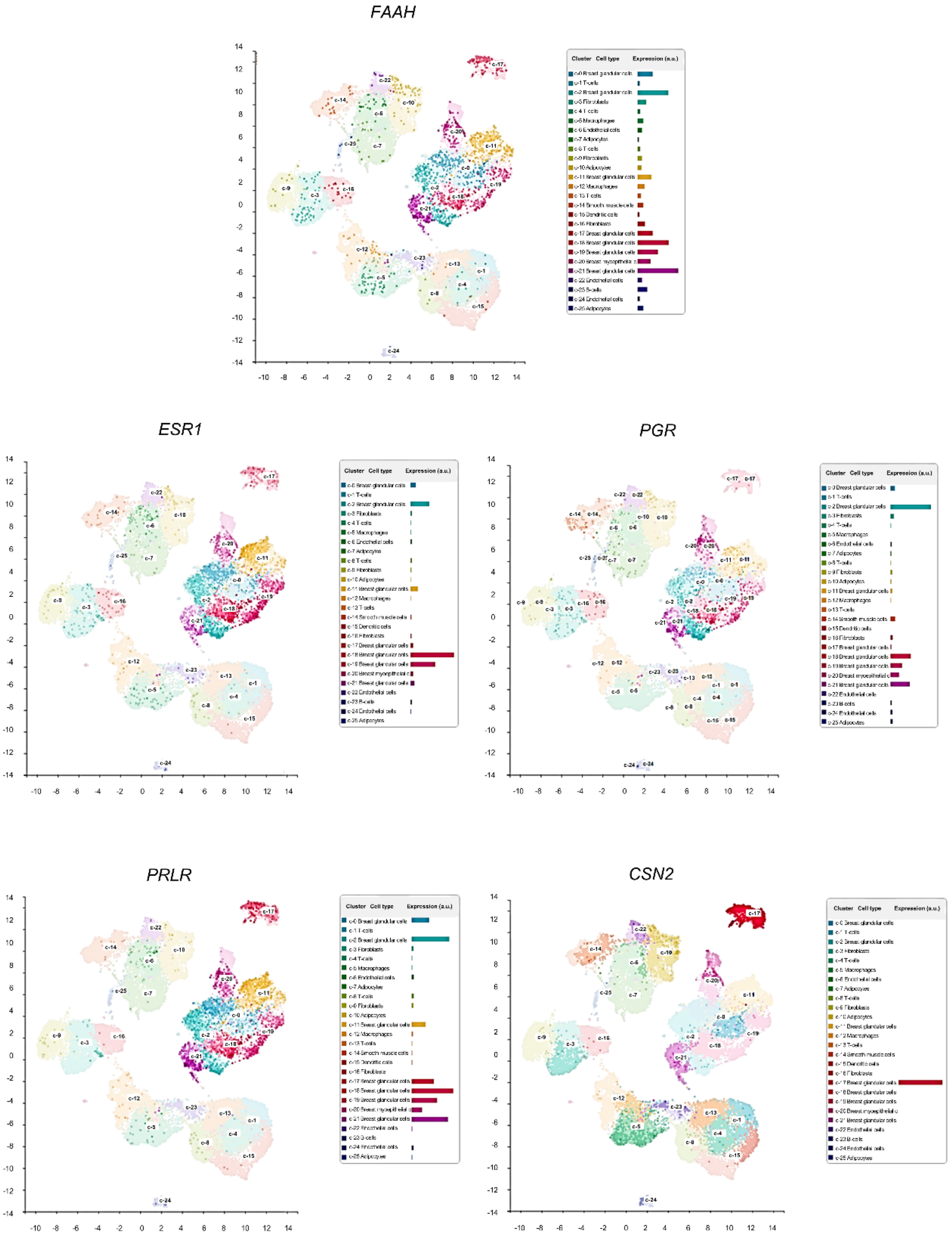

**Supplementary Figure 1. FAAH is expressed in luminal hormone-sensing cell populations.** t-SNEs plots representing mRNA expression of *FAAH* and cell population-specific genes in putative cell clusters of mammary cell populations defined by scRNA-seq analysis of developing mouse mammary glands as published by (Thul et al. 2017). t-SNEs plots are colored by the normalized log-transformed expression of each of the genes. *ESR1*: estrogen receptor alpha; *PGR*: progesterone receptor; *PRLR*: prolactin receptor; *CSN2*: β-Casein.

Supplementary Figure 2

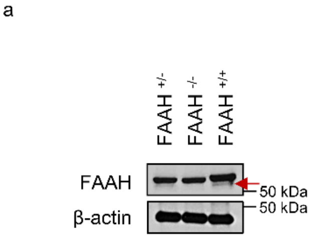


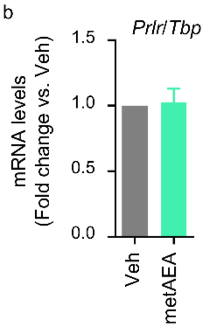

**Supplementary Figure 2. a** Representative WB analysis of FAAH in whole mammary gland lysates from FAAH $^{+/+}$, FAAH $^{+/-}$, and FAAH $^{-/-}$ mice. The WB band pattern in mouse mammary tissue is characterized by a double band where the lower one (pointed with an arrow) is the only which is absent in the FAAH $^{-/-}$ mice. **b** qPCR analysis of *Prlr* mRNA expression in HC11 cells after being differentiated in the presence of the metAEA at 0.5 μM. Expression levels were normalized against *Tbp*. Data are shown as mean ± SEM of n=3 biologically independent experiments. Student's t-test: ns.
